# Cross-disorder genetic analyses implicate dopaminergic signaling as a biological link between Attention-Deficit/Hyperactivity Disorder and obesity measures

**DOI:** 10.1101/573808

**Authors:** Nina Roth Mota, Geert Poelmans, Marieke Klein, Bàrbara Torrico, Noèlia Fernàndez-Castillo, Bru Cormand, Andreas Reif, Barbara Franke, Alejandro Arias Vásquez

## Abstract

Attention-Deficit/Hyperactivity Disorder (ADHD) and obesity are frequently comorbid, genetically correlated, and share brain substrates. The biological mechanisms driving this association are unclear, but candidate systems, like dopaminergic neurotransmission and circadian rhythm, have been suggested. Our aim was to identify the biological mechanisms underpinning the genetic link between ADHD and obesity measures and investigate associations of overlapping genes with brain volumes. We tested the association of dopaminergic and circadian rhythm gene sets with ADHD, body mass index (BMI), and obesity (using GWAS data of N=53,293, N=681,275, and N=98,697, respectively). We then conducted genome-wide ADHD-BMI and ADHD-obesity gene-based meta-analyses, followed by pathway enrichment analyses. Finally, we tested the association of ADHD-BMI overlapping genes with brain volumes (primary GWAS data N=10,720–10,928; replication data N=9,428). The dopaminergic gene set was associated with both ADHD (P=5.81×10^−3^) and BMI (P=1.63×10^−5^), the circadian rhythm was associated with BMI (P=1.28×10^−3^). The genome-wide approach also implicated the dopaminergic system, as the *Dopamine-DARPP32 Feedback in cAMP Signaling* pathway was enriched in both ADHD-BMI and ADHD-obesity results. The ADHD-BMI overlapping genes were associated with putamen volume (P=7.7×10^−3^; replication data P=3.9×10^−2^) – a brain region with volumetric reductions in ADHD and BMI and linked to inhibitory control. Our findings suggest that dopaminergic neurotransmission, partially through DARPP-32-dependent signaling and involving the putamen, is a key player underlying the genetic overlap between ADHD and obesity measures. Uncovering shared etiological factors underlying the frequently observed ADHD-obesity comorbidity may have important implications in terms of prevention and/or efficient treatment of these conditions.

## INTRODUCTION

Attention-Deficit/Hyperactivity Disorder (ADHD) is a psychiatric disorder characterized by developmentally inappropriate and impairing levels of inattention and/or hyperactivity and impulsivity. The prevalence of ADHD is estimated as 5.3% during childhood/adolescence [1] and around 2.8% during adulthood [2]. ADHD is among the most heritable psychiatric disorders, with heritability estimates around 74% [3]. It follows a multifactorial pattern of inheritance, where multiple genetic and environmental factors, each of small effect, and their interplay can contribute to its pathophysiology. A recent genome-wide association study (GWAS) meta-analysis identified the first genome-significant associations for ADHD [4].

High comorbidity rates are a hallmark of ADHD, further increasing disease burden. These comorbidities include both psychiatric and non-psychiatric (somatic) diseases and traits [5]. Among the most frequently reported comorbid somatic conditions in ADHD is obesity [6]. Obesity is nowadays one of the major health problems worldwide, resulting in a large economic burden and significant decrease in life expectancy [7]; its prevalence keeps rising [8]. Obesity is usually classified according to body mass index (BMI), which is calculated as weight in kilograms divided by the height in meters squared (kg/m^2^). A BMI >25 kg/m^2^ signals overweight and a BMI >30 kg/m^2^ is regarded as obesity, which can be further subdivided into classes defined based on increasing BMI [9]. The genetic contribution to obesity and related phenotypes has been extensively studied, and heritability estimates range from 50% up to 90% [10]. Several GWASs have been conducted on obesity and BMI. For BMI, the most recent GWAS meta-analysis included nearly 700,000 individuals and identified 536 associated genomic loci [11]. A previous GWAS on 158,864 participants with BMI information compared normal weight individuals to those with obesity classes I, II, and III [12]. The authors concluded that associations found with categorical phenotypes are highly overlapping with those obtained by using BMI as a quantitative trait [12].

The reported prevalence of ADHD among adults seeking weight loss treatment for obesity is around 27%, reaching up to 43% when considering only those with extreme obesity (i.e. class III) [13, 14]. This rate is over ten times higher than the prevalence of ADHD in adults in the general population [2]. Likewise, two recent meta-analyses show a higher than expected prevalence of overweight and/or obesity in ADHD, both during childhood/adolescence and adulthood, with odds ratios up to 1.55 and strongest effects in adults [15, 16]. Importantly, the association between ADHD and obesity was no longer significant when the analysis was limited to participants receiving pharmacological treatment for ADHD [15].

Specific factors underlying the comorbidity between ADHD and obesity remain largely unknown. Recently, significant genetic correlations between ADHD and BMI (rg=0.21–0.26, [4, 17]) and between ADHD and obesity (ranging from rg=0.285 to rg=0.338 for different obesity classes) and other obesity-related phenotypes have been reported [4]. These findings highlight the involvement of genetic factors in the observed epidemiological overlap between ADHD and obesity measures and provide an entry point for the investigation of specific biological processes involved.

In addition to clinical and genetic overlap between ADHD and obesity measures, volumetric differences in specific brain regions have been associated with ADHD [18] and/or obesity/BMI [19]. In particular, volumes of putamen and nucleus accumbens are reduced in ADHD and are negatively correlated with BMI in the general population [18, 19]. Given that subcortical volumes have also been shown to be heritable traits [20], one may wonder whether shared genetic factors between ADHD and obesity measures could also be associated with volumetric variation in these specific subcortical brain regions.

Some candidate biological systems have been suggested to underly ADHD comorbidity patterns, including dopaminergic neurotransmission and circadian rhythm systems. These two candidate mechanisms have been selected as the main focus of a large European Union consortium aimed at studying comorbid conditions of ADHD (CoCA; https://coca-project.eu/), of which this study is part.

Altered reward processing and impaired inhibitory control, key features of ADHD, are thought to be the outcome of dysregulated dopaminergic neurotransmission [21]. The central role of the dopaminergic system on ADHD is further supported by the dopamine transporter protein being the main target of methylphenidate, the medication of first choice in the pharmacological treatment of ADHD [22]. Studies in humans and animal models have also linked disturbances in dopaminergic neurotransmission and downstream processes to obesity [23, 24]. Overeating may represent an attempt of obese people to compensate for their reduced reward sensitivity [23].

Circadian rhythm-related traits (e.g. eveningness) and disturbances (e.g. sleep problems) have been repeatedly associated with ADHD and/or ADHD symptoms [25]. These problems have also been linked to BMI variation and obesity. Disrupted circadian rhythm signaling may lead to obesity through temporal alterations in eating behavior and changes in metabolic hormones [26]. Two manifestations of circadian rhythm disruption in particular, sleeping problems (i.e. altered sleep duration) and an unstable eating pattern (e.g. skipping breakfast and binge eating later in the day), may mediate the observed association between ADHD symptoms and BMI [27].

In this paper, we aimed to identify shared etiological factors underlying the observed associations of ADHD with obesity measures, and to explore the relationship of overlapping genes with brain volumes. Specifically, we conducted (1) candidate gene-set association analyses and (2) genome-wide gene-based cross-disorder(/trait) meta-analyses, from which the identified overlapping genes were taken forward for (3.1) pathway enrichment analyses and (3.2) testing gene-set association with brain volumes.

## MATERIALS AND METHODS

### Participant samples

This study used publicly available summary statistics of GWAS of ADHD, BMI, obesity, and selected brain volumes. These are briefly described below, and further information is provided in the **Supplementary Material**. These studies had been approved by local ethics committees and had obtained the required informed consents (as described in earlier publications [4, 11, 12, 20, 28]).

The ADHD data was derived from 19,099 cases and 34,194 controls, composed by samples from the Lundbeck Foundation Initiative for Integrative Psychiatric Research (iPSYCH) and the Psychiatric Genomics Consortium (PGC) samples of European ancestry [4].

For BMI, we used summary statistics from the most recent BMI GWAS of European ancestry (N=681,275) from the Genetic Investigation of ANthropometric Traits (GIANT) consortium [29] and UK Biobank [11].

For obesity, summary statistics from a GWAS from European ancestry cohorts within the GIANT consortium on obesity class I were used (N=32,858 cases, N=65,839 controls) [12]. Subjects in that study were considered as cases for obesity class I if they had BMI≥30 kg/m^2^; controls had a BMI<25 kg/m^2^.

Summary statistics for selected brain volumes were derived from GWAS by the Enhancing Neuro Imaging Genetics through Meta-Analysis (ENIGMA) consortium (N range=10,720–10,928)[20]. Furthermore, we used GWAS summary statistics from the UK Biobank samples (N=8,428) [28] for replication of significant findings.

### ADHD – BMI genetic correlation analysis

Due to the large increase in sample size of the most recently published GWAS meta-analysis on BMI, we conducted linkage disequilibrium (LD) score regression analysis [30] to (re-)estimate the genetic correlation between ADHD and BMI using summary statistics of the largest GWASs currently available for each phenotype. We used pre-computed LD scores based on European samples from the 1000 Genomes Project as indicated in https://github.com/bulik/ldsc/wiki/Heritability-and-Genetic-Correlation.

### Hypothesis-driven, candidate gene-set approach

#### Gene-set association analyses

In order to assess the links of dopaminergic neurotransmission and circadian rhythm pathways with ADHD, BMI, and obesity, we assembled gene sets and tested their associations to the individual phenotypes of interest using the GWAS summary statistics described above. Dopaminergic neurotransmission and circadian rhythm gene sets were assembled based on the *Kyoto Encyclopedia of Genes and Genomes (KEGG)* and the *Gene Ontology (GO)* databases. The final dopaminergic (DOPA) and circadian rhythm (CIRCA) gene sets comprised 264 and 284 unique autosomal genes, respectively. Details on the selection of the gene sets are provided in the **Supplementary Material**.

Gene-set association analyses were performed using MAGMA software (version 1.05b [31]). We first carried out single gene-based analyses to assess the degree of association of each gene (i.e. gene-based P-value) with each phenotype. Next, we tested the association of each gene set, through competitive analyses, by aggregating the gene-based P-values according to their presence (or not) in the gene sets (more detailed description in **Supplementary Material**). We used a conservative Bonferroni correction to account for the six gene-set tests (i.e. (DOPA, CIRCA) x ADHD, BMI, obesity); hence, the gene-set significance P-value threshold was set to 8.33×10^−3^.

### Data-driven, genome-wide approach

#### Gene-based cross-disorder/trait meta-analyses

In addition to the hypothesis-driven approach described above, we performed genome-wide gene-based cross-disorder(/trait) meta-analyses by using gene-based P-values for ADHD, BMI, and obesity (obtained as described above) and the gene meta-analysis option in MAGMA software (version 1.05b [31]). The weighted Stouffer’s Z method was used to combine the Z-scores for each gene across cohorts, with weights set to the square root of the sample size each Z-score is based on (i.e. accounting for the fact that sample sizes vary per SNP – and thus per gene – within and between GWAS summary statistics). Since we were interested in the combined effect of each gene on both phenotypes in each pair-wise meta-analysis, only genes present in both gene-based GWAS results were included. The gene-based P-value threshold for genome-wide significance was set to 0.05 divided by the number of genes in each gene-based meta-analysis.

From each pair-wise gene-based cross-disorder(/trait) meta-analysis, the genome-wide significant genes that increased significance by at least one order of magnitude compared to each of the original gene-based results (i.e. P_meta-analysis_ < P_ADHD_/10 and P_meta-analysis_ < P_(obesity or BMI)_/10) were considered as overlapping genes and taken forward for follow-up analyses. This measure was taken in order to avoid including genes for which the association signal is driven solely by one of the phenotypes being meta-analyzed, especially considering the difference in the GWAS sample sizes.

#### Canonical pathway enrichment analyses

The sets of ADHD-BMI and ADHD-obesity overlapping genes were then individually tested for enrichment of canonical pathways using Ingenuity Pathway Analysis (http://www.ingenuity.com; QIAGEN Bioinformatics, Redwood City, California, USA), using its default parameters and Benjamini-Hochberg correction for multiple testing (see **Supplementary Material** for details).

#### Variance explained by ADHD – BMI overlapping genes

We used stratified LD score regression [32] in order to estimate the proportion of the SNP-heritability explained by the set of ADHD-BMI overlapping genes in each of these phenotypes, also testing for heritability enrichment in this set of genes. This variation of partitioned heritability analyses compares the proportion of SNPs (Prop.SNPs) included in the (ADHD-BMI gene set) annotation and the proportion of the SNP-heritability (Prop.h2) accounted for by this to the total number of SNPs and total SNP-heritability. By dividing these two measures (i.e. Prop.h2/Prop.SNPs), an enrichment value and its significance can be calculated, jointly modeling the gene set annotation and the ’baseline model’ of LD score analyses [32].

#### ADHD – BMI overlapping genes and brain volumes

The identified ADHD-BMI overlapping genes were also taken forward in order to test their association, as a gene set, with brain volumes previously found associated with both ADHD [18] and BMI [19] by neuroimaging studies (namely, the putamen and the nucleus accumbens). For these analyses we used GWAS summary statistics of brain volumes from the ENIGMA consortium and the UK Biobank, the latter being used as a replication sample for significant findings (further sample details provided above and in **Supplementary Material**). As exploratory analyses, we also tested the associations of the set of ADHD-BMI overlapping genes with those volumes previously associated with only one of these conditions (i.e. either only ADHD or BMI). The gene-set analyses were carried out in MAGMA software (version 1.05b [31]), in the same manner as described above.

## RESULTS

### ADHD–BMI genetic correlation

The ADHD-BMI genetic correlation was estimated as r_g_=0.3157 (SE=0.0246; P=8×10^−38^). This is similar to estimates based on smaller BMI data sets as well as to estimates for the obesity classes previously reported [4, 17] and mentioned in the introduction.

### DOPA and CIRCA gene-set associations with ADHD, BMI, and obesity

We tested the association of two gene sets – DOPA (264 genes) and CIRCA (284 genes) – with ADHD, BMI, and obesity. Results of these gene-set analyses are shown in **Table 1**. The DOPA gene set was significantly associated with both ADHD (P=5.81×10^−3^) and BMI (P=1.63×10^−5^); the CIRCA gene set was associated with BMI (P=1.28×10^−3^). These results were not driven by one or few individual genes that were highly associated with either ADHD or BMI (**Supplementary Table S1**).

**Table 1.** Gene-set association results of dopaminergic (DOPA) and circadian rhythm (CIRCA) systems with ADHD, BMI, and obesity.

|  | **DOPA<sup>a</sup>** | **CIRCA<sup>b</sup>** |
| --- | --- | --- |
| **ADHD<sup>c</sup>** | **5.81x10<sup>-3</sup>** | 0.521 |
| **BMI<sup>d</sup>** | **1.63x10<sup>-5</sup>** | **1.28x10<sup>-3</sup>** |
| **Obesity<sup>e</sup>** | 0.050 | 0.205 |

Values shown are association P-values. Significant associations are highlighted in bold.
<sup>a</sup>DOPA gene-set analyses are based on 261, 245, and 248 genes from the ADHD, BMI, and obesity GWAS summary statistics, respectively.
<sup>b</sup>CIRCA gene-set analyses are based on 281, 272, and 273 genes from the ADHD, BMI, and obesity GWAS summary statistics, respectively.
<sup>c</sup>European ancestry iPSYCH-PGC ADHD GWAS [4].
<sup>d</sup>GIANT-UK Biobank BMI GWAS [11].
<sup>e</sup>GIANT obesity (class I) GWAS [12].

### ADHD-BMI and ADHD-obesity gene-based meta-analyses

The gene-based cross-disorder(/trait) meta-analysis between ADHD and BMI resulted in 1684 genome-wide significant genes, while the one for ADHD and obesity resulted in 22 significant genes. Of those, 211 genes for the ADHD-BMI meta-analysis and 9 genes for the ADHD-obesity meta-analysis, showed an increase in their association significance (i.e. decrease in P-value) of at least one order of magnitude compared to both individual GWASs. These genes, which were all at least nominally significant in the original GWASs being meta-analyzed, are listed in **Supplementary Tables S2** and **S3**.

**Table 2.** Canonical pathways with significant enrichment in the ADHD-BMI gene-based meta-analysis.

|  | CREB Signaling<br>in Neurons | Synaptic Long<br>Term Depression | Synaptic Long Term<br>Potentiation | Dopamine-DARPP32<br>Feedback in cAMP<br>Signaling |
| --- | --- | --- | --- | --- |
| P-value | 4.11x10<sup>-5</sup> | 5.68x10<sup>-5</sup> | 2.17x10<sup>-4</sup> | 2.19x10<sup>-4</sup> |
| P-value - B-H corrected | 7.95x10<sup>-3</sup> | 7.95x10<sup>-3</sup> | 1.53x10<sup>-2</sup> | 1.53x10<sup>-2</sup> |
| Canonical Pathway size<br>(number of genes) | 212<sup>a</sup> | 188<sup>b</sup> | 127<sup>c</sup> | 165<sup>d</sup> |
| ADHD-BMI genes<sup>e</sup> in<br>the pathway | 10 | 9 | 7 | 8 |
|  | *CACNA1D*<sup>f,g</sup> | *CACNA1D*<sup>f,g</sup> | *CREB3L3*<sup>f</sup> | *CACNA1D*<sup>f,g</sup> |
|  | *CREB3L3*<sup>f</sup> | *GNAT1* | *GRIA1*<sup>f,g</sup> | *CREB3L3*<sup>f</sup> |
|  | *GNAT1* | *GRIA1*<sup>f,g</sup> | *GRM4* | *CSNK1G2* |
|  | *GRIA1*<sup>f,g</sup> | *GRID2* | *ITPR3*<sup>f,g</sup> | *ITPR3*<sup>f,g</sup> |
|  | *GRID2* | *GRM4* | *PLCL1* | *PLCL1* |
|  | *GRIK5* | *IGF1R* | *PPP1R3A* | *PPP1R3A* |
|  | *GRM4* | *ITPR3*<sup>f,g</sup> | *PRKAG1*<sup>g</sup> | *PPP2R3A*<sup>f</sup> |
|  | *ITPR3*<sup>f,g</sup> | *PLCL1* |  | *PRKAG1*<sup>g</sup> |
|  | *PLCL1* | *PPP2R3A* |  |  |
|  | *PRKAG1*<sup>g</sup> |  |  |  |

<sup>a</sup>211 unique genes could be trace back to the NCBI 37.3 gene mapping file, where 207 of them were located in autosomes. The number of nominal genes from this pathway and the number of genes present in the corresponding gene-based results are given as (#Nominal genes/#Genes present): ADHD – 42/203; BMI – 113/194; ADHD-BMI – 120/193. Number of nominal genes from this pathway that are part of the DOPA or CIRCA gene sets: DOPA – 62; CIRCA – 70. Of the 211 genes found in this pathway, 110, 100 and 78 genes overlap with those in the Synaptic Long Term Depression, Synaptic Long Term Potentiation, and Dopamine-DARPP32 Feedback in cAMP Signaling pathways, respectively.
<sup>b</sup>181 genes found in the gene mapping file, 177 of them were autosomes. (#Nominal genes/#Genes present): ADHD – 33/177; BMI – 99/168; ADHD-BMI – 104/168. DOPA – 43; CIRCA – 35. Of the 181 genes found in this pathway, 67 and 61 genes overlap with those in the Synaptic Long Term Potentiation and Dopamine-DARPP32 Feedback in cAMP Signaling pathways, respectively.
<sup>c</sup>125 genes found in the gene mapping file, 123 of them were autosomes. (#Nominal genes/#Genes present): ADHD – 25/122; BMI – 76/113; ADHD-BMI – 79/112. DOPA – 43; CIRCA – 42. Of the 125 genes found in this pathway, 83 genes overlap with those in the Dopamine-DARPP32 Feedback in cAMP Signaling pathway.
<sup>d</sup>156 genes found in the gene mapping file, 155 of them were autosomes. (#Nominal genes/#Genes present): ADHD – 29/154; BMI – 88/143; ADHD-BMI – 91/142. DOPA – 65; CIRCA – 58.
<sup>e</sup> Genes from the ADHD-BMI gene-based meta-analysis results, only considering genome-wide significant (at $P_{\text{threshold}}=2.99 \times 10^{-6}$ ) genes with association P-values lower by at least one order of magnitude in the meta-analysis compared to the gene-based results of both ADHD and BMI individually.
<sup>f</sup> Also part of DOPA in the gene-set analysis.
<sup>g</sup> Also part of CIRCA in the gene-set analysis.

The stratified LD score regression showed that the set of 211 ADHD-BMI overlapping genes explain 9.7% and 3.7% of the SNP-heritability of ADHD and BMI, respectively, yielding significant heritability enrichment (h2_E) in this set of genes (ADHD h2_E=8.671, P=1.915×10-14 and BMI h2_E=3.332, P=5.036×10-11).

### Canonical pathway enrichment analyses

Based on the 211 genes from our ADHD-BMI gene-based meta-analysis, the enrichment analysis identified four significant canonical pathways, as shown in **Table 2**. These were *CREB Signaling in Neurons, Synaptic Long Term Depression, Synaptic Long Term Potentiation,* and *Dopamine-DARPP32 Feedback in cAMP Signaling*. The enrichment analysis for the 9 ADHD-obesity genes also rendered four significant canonical pathways: *GABA Receptor Signaling, Corticotropin Releasing Hormone Signaling, Dopamine-DARPP32 Feedback in cAMP Signaling*, and *Huntington’s Disease Signaling* (**Table 3**).

**Table 3.** Canonical pathways with significant enrichment in the ADHD-obesity gene-based meta-analysis.

|  | **GABA Receptor**<br>**Signaling** | **Corticotropin**<br>**Releasing Hormone**<br>**Signaling** | **Dopamine-DARPP32**<br>**Feedback in cAMP**<br>**Signaling** | **Huntington's**<br>**Disease Signaling** |
| --- | --- | --- | --- | --- |
| P-value | 6.69x10<sup>-4</sup> | 1.45x10<sup>-3</sup> | 2.05 x10<sup>-3</sup> | 4.19 x10<sup>-3</sup> |
| P-value - B-H corrected | 2.81x10<sup>-2</sup> | 2.87x10<sup>-2</sup> | 2.87x10<sup>-2</sup> | 4.40 x10<sup>-2</sup> |
| Canonical Pathway size<br>(Number of genes) | 128 | 143 | 165 | 270 |
| ADHD-obesity genes<sup>a</sup> in<br>the pathway | 2 | 2 | 2 | 2 |
|  | *CACNA1D*<sup>b,c</sup><br>*DNM1* | *BDNF*<br>*CACNA1D*<sup>b,c</sup> | *CACNA1D*<sup>b,c</sup><br>*CSNK1G2* | *BDNF*<br>*DNM1* |

<sup>a</sup> Genes from the ADHD-obesity gene-based meta-analysis results, only considering genome-wide significant genes (at $P_{\text{threshold}}=2.97 \times 10^{-6}$ ) with association P-values lower by at least one order of magnitude in the meta-analysis compared to the gene-based results of both ADHD and obesity individually.
<sup>b</sup> Also part of DOPA in the gene-set analysis.
<sup>c</sup> Also part of CIRCA in the gene-set analysis.

One pathway, the *Dopamine-DARPP32 Feedback in cAMP Signaling*, was found enriched in the two analyses. In total, proteins encoded by eight unique genes derived from our meta-analyses operate in this canonical pathway (**Tables 2 and 3**). Combining the enrichment analysis with a literature search, we constructed a schematic representation of the *Dopamine-DARPP32 Feedback in cAMP Signaling* pathway, which is shown in **Figure 1** and described in detail in the **Supplementary Material**.

**Figure 1.**
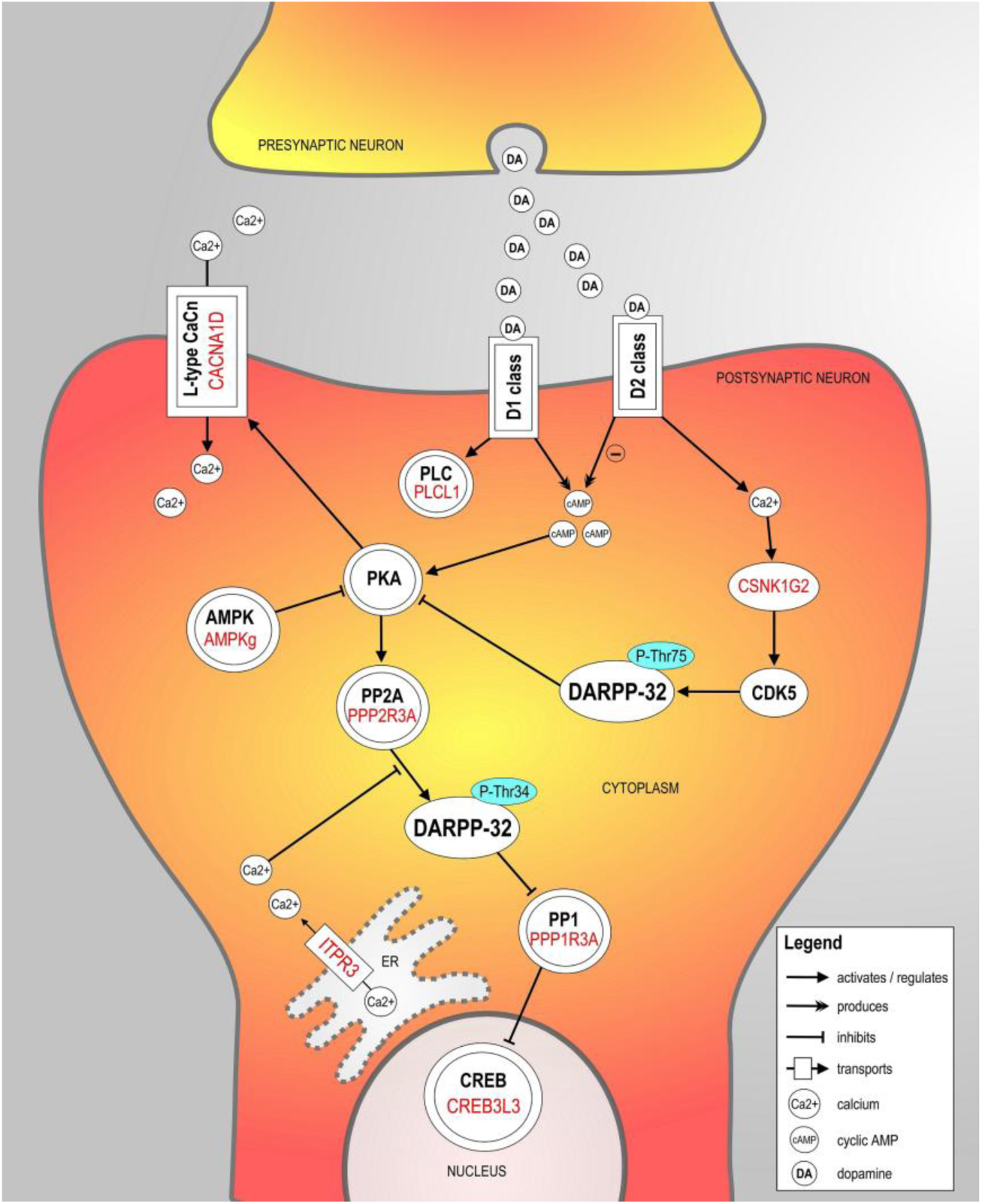
Schematic representation of the *Dopamine-DARPP32 Feedback in cAMP Signaling* pathway. The proteins encoded by the eight genome-wide significant genes derived from the ADHD-BMI gene-based meta-analysis results (Table 2) are contextualized and highlighted in red in the pathway. A detailed description of the pathway in provided in the **Supplementary Material**. For clarity and simplicity, additional proteins in the pathway are omitted. Protein groups or complexes are shown with double margins.

Welcome Trust participants had been included both in the iPSYCH-PGC and the GIANT GWASs; we therefore performed a secondary ADHD-BMI gene-based cross-disorder(/trait) meta-analysis to address the small sample overlap between the data sets (for further information, see **Supplementary Material**). Analysis after excluding those participants from the ADHD GWAS-MA resulted in 202 genes of interest, highly overlapping with the 211 genes from the main ADHD-BMI meta-analysis results (182 overlapping genes), where the *Dopamine-DARPP32 Feedback in cAMP Signaling* pathway remained significantly associated with the phenotype through the canonical pathway enrichment analysis.

### ADHD-BMI overlapping genes and brain volumes

The set of 211 ADHD-BMI overlapping genes was also used to test the association with volumetric variation of brain structures. As shown in **Table 4**, the ADHD-BMI gene set was significantly associated with putamen volume in the ENIGMA GWAS-MA (P=7.7×10^−3^). This association was replicated in the UK Biobank GWAS results (P=3.9×10^−2^). Associations with other subcortical volumes and intracranial volume were also tested, as exploratory analyses, and were all non-significant (**Table 4**).

**Table 4.** Gene-set association results of the set of 211 ADHD-BMI overlapping genes with brain volumes.

| Brain volume | ENIGMA GWAS<sup>a</sup> |  | UK Biobank GWAS<sup>b</sup> |
| --- | --- | --- | --- |
|  | Mean N | P-value | P-value |
| *Main analyses<sup>c</sup>* |  |  |  |
| Putamen | 10,829 | **7.65x10<sup>-3</sup>** | **3.94x10<sup>-2</sup>** |
| Nucleus accumbens | 10,887 | 0.517 | - |
| *Exploratory analyses<sup>d</sup>* |  |  |  |
| Amygdala | 10,928 | 0.235 |  |
| Caudate | 10,914 | 0.114 |  |
| Hippocampus | 10,845 | 0.714 |  |
| Pallidum | 10,829 | 0.975 |  |
| Intracranial | 10,720 | 0.470 |  |

Significant associations are highlighted in bold.
<sup>a</sup>GWAS summary statistics from the ENIGMA consortium, as described by Hibar et al. [20]. Previous to gene-set analyses, the NeuroIMAGE cohort (N=154), which includes ADHD cases, was removed from the ENIGMA data.
<sup>b</sup>Replication sample - GWAS summary statistics from the UK Biobank cohort, including N=8,428 individuals, as described by [28].
<sup>c</sup>Brain volumes previously associated with both ADHD [18] and BMI [19].
<sup>d</sup>Brain volumes associated only with ADHD or BMI [18, 19].

## DISCUSSION

In this paper, we aimed to uncover biological mechanisms underlying the observed genetic associations between ADHD and obesity measures. Based on known and self-derived genetic correlation estimates for ADHD and BMI/obesity obtained from the world-wide largest data sets for each phenotype, we first applied a hypothesis-driven testing approach of two selected gene sets (DOPA and CIRCA), which showed that the dopaminergic neurotransmission system partially explains the genetic overlap between ADHD and BMI. Our data-driven, genome-wide approach subsequently showed that dopaminergic signaling, specifically *Dopamine-DARPP32 Feedback in cAMP Signaling*, was significantly enriched in both the ADHD-BMI and the ADHD-obesity gene-based meta-analysis results.

Both ADHD and obesity measures have been linked to disturbances in dopaminergic signaling. Alterations of the brain’s executive and reward circuits – modulated by mesocortical and mesolimbic dopamine, respectively – have been postulated as the basis of the deficient inhibitory control and impaired reward processing characteristics of ADHD [21]. The ability to resist the impulse to eat desirable foods, and an appropriate reward-response to those, also require proper functioning of these dopamine-regulated processes [23, 24]. For example, impulsive eating, as a result of a high arousal response to a potential reward and impaired inhibitory control, can lead to weight gain and obesity [33]. Eating behavior is also dependent on the hypothalamic homeostatic system, which comprises hormonal regulators of energy balance – such as insulin, leptin, and gut hormones – and controls hunger, satiety, and adiposity [23]. Increasing evidence suggests that such metabolic hormones also affect food-related sensitivity of the dopaminergic reward system [34], pointing to an overlap between the homeostatic and reward/reinforcement systems related to obesity [23].

Also confirming our hypothesis, the CIRCA gene set was associated with BMI, but the absence of a significant association with ADHD was unexpected. ADHD has previously been associated with altered circadian rhythmicity at molecular, endocrine, and behavior levels [35]. Furthermore, zebrafish mutants for *per1b*, a key gene in circadian rhythm regulation, and *Per1*-knockout mice display hyperactive, impulsivity-like, and attention deficit-like behaviors [36]. The lack of a significant association between ADHD and the CIRCA gene set in our study may be due to a true lack of effect of the circadian rhythm pathway on ADHD. However, given that some of the CIRCA genes are among the cross-disorder(/trait) overlapping genes, it is also possible that there is a true (unobserved) effect but that the gene set we assembled was not appropriate/informative enough to detect such association.

Going beyond candidate gene-set analyses, we conducted data-driven, genome-wide ADHD-BMI and ADHD-obesity gene-based meta-analyses. Cross-disorder(/trait) overlapping genes were carried forward into two follow-up approaches: one testing the association of (ADHD-BMI) overlapping genes with specific subcortical brain volumes previously linked to these phenotypes and the other aimed at identifying enriched biological pathways underlying the shared heritability. Both follow-up approaches again pointed to a role of the dopaminergic system. Through the first, we observed a significant association of ADHD-BMI overlapping genes with putamen volume in two independent samples (**Table 4**). This finding is of particular interest given the strong role of the dopaminergic system in this brain region and the prominent involvement of the putamen in inhibitory control functioning, one of the key neurobiological features suggested to be altered both in ADHD and obesity [18, 19]. The second follow-up approach showed several pathways significantly enrichment in the ADHD-BMI and in the ADHD-obesity results. Dopamine signaling was at the heart of the pathway that was significantly enriched in both analyses, i.e. the *Dopamine-DARPP32 Feedback in cAMP Signaling* pathway. This postsynaptic pathway centers around the Dopamine- and cAMP-regulated neuronal phosphoprotein (DARPP-32; also known as Protein phosphatase 1 regulatory subunit 1B (PPP1R1B)), the phosphorylation state of which modulates dopaminergic neurotransmission (see **Figure 1** and description in the **Supplementary Material** for details).

DARPP-32 is primarily expressed in postsynaptic dopaminergic neurons in the dorsal striatum (i.e. brain structure which includes, in addition to the caudate, the putamen; see results above for the association of ADHD-BMI overlapping genes and brain volumes), which is involved in certain executive functions, such as inhibitory control, and in the ventral striatum, which is the main brain region responsible for reward processing (https://gtexportal.org/home/gene/PPP1R1B). As described above, poor inhibitory control and altered reward processing, in the form of steeper delay discounting, are key neurobiological circuitries implicated in both ADHD and obesity [21, 23]. Further evidence linking dopamine-DARPP-32 signaling, reward processing and the brain comes from findings in animal models. Upon investigation of the consequences of frustrated expected reward of palatable food on gene expression in the mouse brain, *Dopamine-DARPP32 Feedback in cAMP Signaling* pathway was found to be enriched among differentially expressed genes, both the ventral striatum and in frontal cortex [37].

DARPP-32 modulates the effects of dopamine on cAMP/PKA-dependent gene transcription through transcription factors of the cyclic AMP-responsive element-binding (CREB) complex (**Figure 1**), and CREB dysregulation has been linked to both ADHD [38] and obesity [39]. Of note, the *CREB Signaling in Neurons* pathway was also significantly enriched in our ADHD-BMI gene-based meta-analysis, along with two other partially overlapping pathways involved in synaptic plasticity processes (namely, the *Synaptic Long Term Depression* and the *Synaptic Long Term Potentiation* pathways; **Table 2**), which are also closely related to dopamine DARPP-32 signaling.

Additional evidence for an involvement of DARPP-32 signaling to the ADHD-BMI/obesity overlap comes from the study of rare variants. The most common form of monogenic obesity is caused by mutations in the melanocortin 4 receptor (*MC4R*) gene [40], and MRC4 signaling is known to activate DARPP-32 [41]. In addition to early-onset obesity, a higher prevalence of ADHD has been reported in *MC4R* mutation-carriers [42]. It has been hypothesized that such co-occurrence may be, in part, underpinned by reward processing deficits [43], and animal studies provide further support regarding the involvement of MC4R signaling and dopaminergic-dependent reward processing [41].

Our study has strengths and limitations. A clear strength is that we make use of the largest GWAS results available for each of the phenotypes being investigated. The sample sizes used to generate the (European ancestry) summary statistics used here were, in total, more than 53,000 for the iPSYCH-PGC ADHD GWAS, up to 700,000 for the GIANT-UK Biobank BMI GWAS, and almost 99,000 for the GIANT obesity GWAS. Obesity measures were, therefore, assessed both as a trait and a state. Although we performed the (categorical) obesity analysis using GWAS data from the obesity class with the largest sample size (obesity class I Ncases= 32,858 Ncontrols=65,839; class II Ncases=9,889 Ncontrols=62,657; class III Ncases=2,896 Ncontrols=47,468; [12]), it is possible that the quantitative nature of BMI and the much larger sample size of the BMI GWAS provide more powerful analyses/results than with the obesity GWAS class I, which may account, at least in part, for some of the differences observed between the BMI and Obesity results. All GWAS summary statistics used here are derived from individuals with European ancestry; the homogeneous background can be a strength given that genetic analyses can be sensible to population stratification, but we also would like to highlight the need of large studies on more diverse populations. Another strength is that we did not restrict our gene set assembly to single GO-terms or KEGG pathways, but applied a more inclusive approach regarding the processes involved. For dopaminergic neurotransmission, we thus assembled a gene set (DOPA) that was subsequently found to be significantly associated with ADHD and BMI. This contrasts with the approach adopted in the iPSYCH-PGC ADHD GWAS paper, which tested dopaminergic candidate genes and GO-term pathways only individually, failing to detect significant associations with ADHD [4]. The large difference in sample sizes between the phenotypes imposed some difficulties when analyzing them together. We minimized such limitations by carrying out gene-based cross-disorder(/trait) meta-analyses in MAGMA, which allows sample sizes to vary between and within samples and accounts for such variation by weighting the effects accordingly. We also opted for performing gene-based – rather than SNP-based – cross-disorder(/trait) meta-analyses. Apart from assuming that the (combined effect of SNPs within) genes represent entities closer to the biological mechanisms, this approach has a reduced statistical burden compared to SNP-based analyses and seems most suitable for these data given the difference in SNP density between the ADHD and the BMI and obesity GWASs (the later ones being restricted to about 2.4 million SNPs present in HapMap 2). An additional advantage of using gene-based approach when meta-analyzing different phenotypes is that it doesn’t rely on *a priori* expectations of concordance of the direction of effects, which avoids information on loci with discordant direction of effects from being lost. Another limitation we addressed was the presence of overlapping samples, since Welcome Trust participants had been included both in the iPSYCH-PGC ADHD GWAS and the GIANT BMI and obesity GWASs. The reduction in sample size reduced power of our analysis, but findings from the canonical pathway enrichment analysis remained stable. Finally, despite the undeniable genetic component of these complex disorders/traits, the current available sample sizes and techniques applied in genome-wide studies still only allow for a small proportion of the phenotypic variance to be accounted for by common variants genome-wide. However, we strongly believe that identifying the biological pathways shared between disorders represents a promising way forward to a better understanding of comorbidity, which goes far beyond the observed effect sizes of specific genes/pathways and their variance explained. Given the limitations stated above, our results should be interpreted with caution and considered as exploratory until more adequately powered samples and methods are available.

Overall, the findings of the present study identify dopaminergic neurotransmission as a key player underlying the shared heritability of ADHD and BMI/obesity, implicating mechanisms involving DARPP-32 signaling in particular and possibly involving neurobiological features related to putamen, such as inhibitory control. This is especially interesting since DARPP-32 has been directly implicated in the mechanism of action of ADHD medication [44], which has been suggested to attenuate the increased risk for obesity in people with ADHD [15]. The fact that we observe a convergence between the results from hypothesis-driven and hypothesis-free approaches provides extra support to the robustness of our findings. Uncovering critical aspects of the shared etiology underlying the prevalent ADHD-obesity comorbidity may have important implications for clinical outcome, preventive interventions, and/or efficient treatment of these conditions.

