## Supplementary Material for "Cross-disorder genetic analyses implicate dopaminergic signaling as a biological link between Attention-Deficit/Hyperactivity Disorder and obesity measures"

### Supportive information – Methods

#### GWAS summary statistics

The ADHD GWAS summary summary statistics used in our analyses are deriverd from iPSYCH-PGC samples, as described in details by Demontis et al. (1), and can be obtained from the PGC website (https://www.med.unc.edu/pgc/results-and-downloads). Data was filtered to include only SNPs with minor allele frequency (MAF)>0.01 and an imputation quality (INFO)>0.8, totalizing 8,094,094 variants.

The BMI GWAS summary statistics were provided by the GIANT consortium and downloaded from https://portals.broadinstitute.org/collaboration/giant/index.php/GIANT_consortium_data_files. Detailed description of samples, data and analysis procedures can be found in Yengo et al. (2). After downloading, the data was subsequently filtered in order to include only SNPs with MAF>0.01 in our analyses, which yielded a total of 2,336,056 SNPs.

We also used the obsesity (class I) GWAS from the GIANT consortium, which has been described in details by Berndt et al. (3). Data was downloaded from https://portals.broadinstitute.org/collaboration/giant/index.php/GIANT_consortium_data_files and subsequently filtered to include only SNPs with MAF>0.01, resulting in 2,353,324 SNPs.

From the ENIGMA consortium GWAS, detailed description of samples and GWAS data can be found in the original publication by Hibar et al. (4) and summary statistics are available upon request via the ENIGMA website (http://enigma.ini.usc.edu/download-enigma-gwas-results/). The GWASs of subcortical brain volumes were conducted adjusting for intracranial volume. The NeuroIMAGE cohort (N=154), which includes ADHD patients, was removed from the ENIGMA data prior to analyses of the current study. Only SNPs with imputation score of RSQ>0.5 are included in the ENIGMA summary statistics and we further filtered SNPs to include only those with MAF>0.01. The final number of SNPs ranged from 8,404,06 to 8,445,488 for the different brain volumes analysed (i.e. putamen, nucleus accumbens, caudate, pallidum, amygdale, hippocampus and intracranial volume).

GWAS results from brain volumes from the UK Biobank cohort were used as replication data for any significant association found with the ENIGMA data. Detail information of these summary statistics, which include only SNPs with INFO>0.3, are provided by Elliot et al (5). Analyses were restricted to SNPS with MAF>0.01, totalizing 7,723,976 SNPs.

**Candidate gene-set association analyses**

*Gene-set assembly*

For our candidate gene-set analyses, we assembled dopaminergic neurotransmission (DOPA) and circadian rhythm (CIRCA) gene-sets based on the *Kyoto Encyclopedia of Genes and Genomes (KEGG)* and the *Gene Ontology (GO)* databases, queried in September 2016.

DOPA gene set: From the KEGG database, we included genes covered by the Dopaminergic synapse (hsa04728) and/or Tyrosine metabolism (hsa00350) pathways and from the GO database we included genes present in (at least) one of the following terms (GO: accession number): dopamine transport (GO:0015872), dopamine receptor signaling pathway (GO:0007212), dopamine receptor binding (GO:0050780), synaptic transmission, dopaminergic (GO:0001963), dopaminergic neuron axon guidance (GO:0036514), dopaminergic neuron differentiation (GO:0071542), response to dopamine (GO:1903350), and dopamine metabolic process (GO:0042417).

This resulted in 155 genes from KEGG database and 144 genes from GO, of which 24 were in common, totalizing 275 genes. From these, we found 273 genes in the MAGMA gene template ("*KIF5C*" and "*PPP2R3B*" were not present), including 9 genes located on chromosome X, which was not covered by our analyses.

CIRCA gene set: From the KEGG database, we included genes comprised by the Circadian rhythm (hsa04710) and/or Circadian entrainment (hsa04713) pathways and, from the GO database we included genes present in the circadian rhythm (GO:0007623) GO term.

This resulted in 123 genes from the KEGG database and 197 genes from GO, of which 27 were in common, totalizing 293 genes. From these, we found 290 genes in the MAGMA software gene reference template (“*H0Y8X5*”, “*LOC400927-CSNK1E*”, and “*Q59FM5*” were not present), including 6 genes located on chromosome X, which was not covered by our analyses.

*Gene-set association analyses*

The gene-set association analyses were performed using MAGMA software (version 1.05b; (6)). For such, SNPs were annotated to protein-coding genes, according to the location of their transcribed regions in the

Human Genome Build 37, using the NCBI 37.3 gene reference template definitions provided by MAGMA (https://ctg.cncr.nl/software/magma). Genes were considered as present in the GWAS summary statistics being analyzed if they contained at least one SNP located within their transcribed region. Gene-based P-values were then calculated using the SNP-wise mean model (default for summary statistics analysis), which combines the effects of SNPs within a gene and uses the sum of –log(SNP P-value) as test statistic. In order to account for linkage disequilibrium (LD) between the SNPs, we used the European dataset of the 1000 Genomes Phase 3 as reference, provided at MAGMA's website (https://ctg.cncr.nl/software/MAGMA/ref_data/g1000_eur.zip).

We then performed competitive gene-set analyses, which tests whether each gene-set is differently associated with the phenotype compared to the remaining genes on the genome, while taking into account gene size, gene density and LD between genes.

**Canonical pathway enrichment analysis**

The sets of ADHD-BMI and ADHD-obesity overlapping genes, derived from the gene-based cross-disorder(/trait) meta-analyses, were tested for enrichment of canonical pathways using Ingenuity Pathway Analysis (http://www.ingenuity.com; QIAGEN Bioinformatics, Redwood City, California, USA). The Ingenuity Knowledge Base, which incorporates experimental data from published literature as well as data from many other sources, including gene expression and gene annotation databases, currently comprises 692 curated canonical pathways of functionally related and interacting genes/proteins (http://qiagen.force.com/KnowledgeBase/articles/Basic_Technical_Q_A/Available-Canonical-Pathways). For each of these pathways, Ingenuity calculates enrichment P-values using the right-tailed Fisher's exact test and taking into consideration both the total number of genes from the analyzed data set (i.e. the 211 ADHD-BMI or the 9 ADHD-obesity overlapping genes) and the total number of genes that is in the pathway in question according to the Ingenuity Knowledge Base. To account for multiple testing, the enrichment P-value of each pathway is adjusted using Benjamini-Hochberg correction, and only significantly enriched canonical pathways were reported.

### Supportive information – Results

**Secondary ADHD-BMI gene-based meta-analysis: no sample overlap**

Since Welcome Trust participants had been included both in the iPSYCH-PGC (1) and GIANT (2, 3) studies, we performed a secondary analysis to make sure that our results were not driven by this overlap. As described by Demontis *et al.* (1), the ADHD PGC sample from Cardiff is composed of 721 ADHD cases and 5081 Welcome Trust controls. Therefore, we performed a secondary ADHD-BMI gene-based meta-analysis using the leave-one-out summary statistics from the European ancestry iPSYCH-PGC ADHD GWAS without the Cardiff sample (~11% reduction in sample size). Unfortunately, it was not feasible to remove these overlapping samples from the GIANT-UK Biobank BMI GWAS. The canonical pathway enrichment findings from this secondary meta-analysis were reassuring since the *Dopamine-DARPP32 Feedback in cAMP Signaling* pathway remained significantly associated, with seven genes in the pathway and surviving B-H correction ($P = 8.90 \times 10^{-4}$, $P_{B\text{-}H} = 4.57 \times 10^{-2}$). Since our secondary results do not show a substantial bias due to sample overlap, we opted for presenting the analysis with the largest sample size as the main analysis.

**Detailed description of Figure 1**

In **Figure 1,** we show a schematic representation of the *Dopamine-DARPP32 Feedback in cAMP Signaling* pathway. In the description below, we contextualize and highlight in bold the proteins encoded by the eight genes derived from our ADHD-BMI gene-based meta-analysis results that are present in this pathway (highlighted in red in **Figure 1**).

Signaling in the *Dopamine-DARPP32 Feedback in cAMP Signaling* pathway starts at the postsynaptic membrane where dopamine released by midbrain neurons binds and activates two types of dopamine receptors, i.e. the D1 class (DRD1 and DRD5) and D2 class (DRD2, DRD3, and DRD4) of receptors (D1 class and D2 class in **Figure 1**) (7). Subsequently, the D1 and D2 receptors work through different G protein alpha subunits to activate and inhibit the enzyme adenylate cyclase, which results in more and less cyclic AMP (cAMP) being synthesized, respectively (7). cAMP in turn activates protein kinase A (PKA), which is inhibited by the kinase AMPK (8, 9) of which the **AMPKg** subunit is encoded by the *PRKAG1* gene (10). Activated PKA has a large number of downstream effects, including regulating the activity of L-type calcium channels (11) – of which **CACNA1D** is a subunit (10) – that are voltage-sensitive and stimulate the entry of calcium ions into neurons (12). Dopamine binding to D2 receptors also leads to a signaling cascade that results in an increased intraneuronal concentration of calcium (13). In addition, activated PKA is involved in activating transcription factors of the cAMP-dependent CREB complex (14, 15) such as **CREB3L3** (10). The CREB complex transcription factors are inhibited by the phosphatase PP1 (16) – which contains the regulatory subunit **PPP1R3A** - leading to a disruption of dopamine-induced and cAMP/PKA-dependent gene transcription, which may ultimately result in various neuropsychiatric disorders - including ADHD (17) - and somatic disorders such as obesity (18, 19). Dopamine binding and activating D1 receptors also leads to the activation of phospholipase C (PLC) enzymes (11, 20) such as **PLCL1** that binds and functionally interacts with PP1 (interaction not shown in Figure 1) (21). The above described dopamine-induced gene transcription is partially controlled by the protein 'Dopamine and cAMP Regulated Phosphoprotein-32' or DARPP-32, through a feedback loop on the cAMP/PKA cascade. Activated PKA stimulates the phosphatase PP2A - of which **PPP2R3A** is a regulatory subunit (10) - that phosphorylates DARPP-32 at Threonine(Thr)34, which makes DARPP-32 a potent inhibitor of PP1 (16). Conversely, an increase of intracellular calcium - through being pumped into the neuron by L-type calcium channels such as **CACNA1D**, downstream of dopamine binding to D2 receptors (see above) and/or trough being released from the endoplasmic reticulum (ER), a process that is mediated by the ER membrane-receptor **ITPR3 (22)** - leads to a dephosphorylation of DARPP-32 at Thr34 (23), rendering the protein inactive again. Interestingly, DARPP-32 can also be phosphorylated at Thr75 by CDK5 (16)- a kinase that itself is activated by CK1 (24), of which **CSNK1G2** is an isoform (10) - which converts DARPP-32 into an inhibitor of PKA (16). In this way, DARPP-32 is a dual-function protein that, depending on where it is phosphorylated, can act either as an inhibitor of PP1 or of PKA, which makes DARPP-32 a critical regulator of dopamine/cAMP/PKA signaling.

**Supplementary Table S1.** Summary of gene-based association results of the ADHD, BMI, and obesity individual GWASs, as well as of the ADHD-BMI and ADHD-obesity gene-based meta-analyses, displaying the number of genome-wide significant (GWsig) genes and if they are included in the candidate DOPA and/or CIRCA gene sets:

| | GWsig genes | Presence of GWsig genes in gene sets: | | |
|---|---|---|---|---|
| | | **DOPA** | **CIRCA** | **DOPA and CIRCA** |
| ADHD[a] | 20 | . | . | . |
| BMI[b] | 1747 | 41 (*ADCY5, ADCY6, AKT3, ARNTL, CACNA1C, CACNA1D, CALML6, COMT, CREB1, CREB3L1, CREB3L3, CTNNB1, DDC, DNM1, DNM2, DRD2, GNAI3, GNAQ, GNAS, GNB1, GNG7, GRIA1, GRIN2A, GSK3B, ITPR3, KLF16, LMX1B, OPRK1, OPRM1, PARK2, PLCB1, PLCB4, PPP1CB, PPP2R3A, PPP2R5C, PPP3CA, PTGS2, RAC1, SCN1A, SLC22A3, SLC6A4*) | 49 (*ADCY3, ADCY5, ADCY6, ADCY9, ADK, AHR, ARNTL, BTBD9, BTRC, CACNA1C, CACNA1D, CALML6, CREB1, CRTC1, DDC, DRD2, EP300, GNAI3, GNAQ, GNAS, GNB1, GNG7, GRIA1, GRIN2A, GSK3B, GUCY1A2, HCRTR1, HCRTR2, ITPR3, LGR4, NCOR1, NFIL3, NLGN1, NOS1AP, NR1H3, NTRK2, OPRL1, PLCB1, PLCB4, PPARG, PPARGC1A, PPP1CB, PRKG1, RORA, RPS6KA5, SLC6A4, SYNCRIP, TP53, ZFHX3*) | 22 (*ADCY5, ADCY6, ARNTL, CACNA1C, CACNA1D, CALML6, CREB1, DDC, DRD2, GNAI3, GNAQ, GNAS, GNB1, GNG7, GRIA1, GRIN2A, GSK3B, ITPR3, PLCB1, PLCB4, PPP1CB, SLC6A4*) |
| Obesity[c] | 26 | . | 1 (*ADCY3*) | . |
| ADHD-BMI[d] meta-analysis | 211 | 8 (*CACNA1D, CREB3L1, CREB3L3, CTNNB1, DNM1, GRIA1, ITPR3, PPP2R3A*) | 7 (*CACNA1D, GRIA1, ITPR3, PRKAG1, RPS6KA5, SYNCRIP, ZFHX3*) | 3 (*CACNA1D, GRIA1, ITPR3*) |
| ADHD-Obesity[e] meta-analysis | 9 | 2 (*CACNA1D, DNM1*) | 1 (*CACNA1D*) | 1 (*CACNA1D*) |

[a.] European ancestry iPSYCH-PGC ADHD GWAS (1).
[b]. GIANT-UK Biobank BMI GWAS (2).
[c] GIANT Obesity (class I) GWAS (3).
[d,e] Meta-analysis of [d.]ADHD-BMI and [e.]ADHD-obesity gene-based results. Only genome-wide significant genes are shown with association P-values lower by at least one order of magnitude in the meta-analysis compared to both the ADHD and the [d.]BMI or [e.]obesity results individually.

**Supplementary Table S2.** Results of the ADHD-BMI gene-based meta-analysis showing the 211 genome-wide significant genes that increased significance in the meta-analysis by at least one order of magnitude compared to both individual GWASs.

| GENE | CHR | START | STOP | ADHD gene-based results | BMI gene-based results | ADHD-BMI gene-based meta-analysis |
|---|---|---|---|---|---|---|
| *SEMA3F* | 3 | 50192562 | 50226508 | 1.83X10-04 | 2.27X10-24 | 1.70X10-26 |
| *FBXL17* | 5 | 107194734 | 107718080 | 4.08X10-05 | 3.17X10-23 | 2.52X10-26 |
| *XKR6* | 8 | 10753654 | 11058875 | 1.06X10-02 | 2.61X10-21 | 1.84X10-22 |
| *GNAT1* | 3 | 50229043 | 50235129 | 1.90X10-03 | 1.40X10-20 | 2.14X10-22 |
| *MAML3* | 4 | 140637545 | 141075233 | 4.81X10-04 | 7.33X10-20 | 4.04X10-22 |
| *CCDC171* | 9 | 15552872 | 15971897 | 3.88X10-03 | 4.14X10-20 | 1.16X10-21 |
| *SOX5* | 12 | 23682438 | 24715383 | 9.20X10-04 | 1.06X10-18 | 9.86X10-21 |
| *PCDH7* | 4 | 30721951 | 31148423 | 1.10X10-03 | 7.55X10-18 | 8.31X10-20 |
| *GATA4* | 8 | 11534433 | 11617510 | 7.44X10-03 | 4.70X10-18 | 2.34X10-19 |
| *SEMA6D* | 15 | 47476403 | 48066420 | 3.11X10-10 | 2.34X10-14 | 2.42X10-19 |
| *TENM2* | 5 | 166406083 | 167691162 | 1.13X10-02 | 1.36X10-17 | 9.78X10-19 |
| *CALN1* | 7 | 71244476 | 71912136 | 4.48X10-05 | 3.51X10-15 | 6.42X10-18 |
| *CACNA1D* | 3 | 53529076 | 53847179 | 3.60X10-04 | 6.19X10-15 | 3.96X10-17 |
| *TRAF3* | 14 | 103243816 | 103377837 | 3.15X10-03 | 1.53X10-15 | 4.01X10-17 |
| *NLRC3* | 16 | 3589033 | 3627392 | 1.02X10-02 | 9.63X10-16 | 6.38X10-17 |
| *STAG1* | 3 | 136055077 | 136471245 | 3.30X10-03 | 3.02X10-15 | 8.34X10-17 |
| *CSMD2* | 1 | 33979609 | 34631443 | 9.04X10-03 | 3.41X10-15 | 2.04X10-16 |
| *ADARB1* | 21 | 46494493 | 46646478 | 2.00X10-03 | 1.86X10-14 | 3.67X10-16 |
| *RSRC1* | 3 | 157823690 | 158262624 | 4.44X10-04 | 5.13X10-14 | 3.94X10-16 |
| *PCCB* | 3 | 135969167 | 136056737 | 2.48X10-03 | 3.94X10-14 | 9.39X10-16 |
| *MSL2* | 3 | 135867760 | 135915522 | 2.22X10-04 | 1.73X10-13 | 9.87X10-16 |
| *MSI2* | 17 | 55333931 | 55762050 | 1.30X10-03 | 9.43X10-14 | 1.48X10-15 |
| *TRAIP* | 3 | 49866028 | 49893992 | 1.14X10-04 | 4.02X10-13 | 1.61X10-15 |
| *IP6K1* | 3 | 49761728 | 49823973 | 1.54X10-04 | 3.68X10-13 | 1.71X10-15 |
| *CAMKMT* | 2 | 44589043 | 44999731 | 1.63X10-04 | 3.86X10-13 | 1.86X10-15 |
| *IGF1R* | 15 | 99192272 | 99507759 | 9.53X10-04 | 1.46X10-13 | 1.94X10-15 |
| *PPL* | 16 | 4932508 | 4987136 | 8.00X10-03 | 3.58X10-14 | 2.00X10-15 |
| *PDE1C* | 7 | 31791666 | 32339016 | 3.63X10-03 | 7.83X10-14 | 2.44X10-15 |
| *TEX29* | 13 | 111973015 | 111996594 | 9.50X10-03 | 4.32X10-14 | 2.73X10-15 |
| *DNASE1* | 16 | 3661772 | 3712689 | 6.14X10-03 | 6.74X10-14 | 3.08X10-15 |
| *ABHD17C* | 15 | 80987635 | 81047962 | 9.52X10-05 | 8.36X10-13 | 3.14X10-15 |
| *PTBP2* | 1 | 97187161 | 97280605 | 2.52X10-03 | 1.45X10-13 | 3.62X10-15 |
| *MST1* | 3 | 49721380 | 49726196 | 3.96X10-04 | 6.71X10-13 | 5.45X10-15 |
| *BSN* | 3 | 49591922 | 49708982 | 1.12X10-05 | 3.85X10-12 | 5.61X10-15 |
| *RNF123* | 3 | 49726950 | 49758962 | 1.03X10-03 | 4.93X10-13 | 7.21X10-15 |
| *ARHGEF7* | 13 | 111767624 | 111958081 | 1.87X10-04 | 1.30X10-12 | 7.22X10-15 |
| *PCDH9* | 13 | 66876966 | 67804468 | 1.74X10-03 | 3.79X10-13 | 7.48X10-15 |
| *MEF2C* | 5 | 88014058 | 88199922 | 3.27X10-08 | 4.66X10-11 | 7.80X10-15 |
| *TSHZ2* | 20 | 51588946 | 52111869 | 1.36X10-02 | 1.61X10-13 | 1.37X10-14 |
| *DNM1* | 9 | 130965634 | 131017528 | 2.46X10-04 | 2.48X10-12 | 1.76X10-14 |

| | | | | | | |
|---|---|---|---|---|---|---|
| *TNKS* | 8 | 9412756 | 9639856 | 1.58X10-02 | 1.99X10-13 | 1.91X10-14 |
| *UBN1* | 16 | 4896666 | 4932363 | 4.90X10-03 | 4.93X10-13 | 1.99X10-14 |
| *CLUAP1* | 16 | 3550945 | 3589048 | 3.93X10-03 | 6.06X10-13 | 2.10X10-14 |
| *UBA7* | 3 | 49842638 | 49851391 | 9.40X10-05 | 5.60X10-12 | 2.32X10-14 |
| *PURG* | 8 | 30853320 | 30891231 | 3.14X10-03 | 8.72X10-13 | 2.68X10-14 |
| *RPS6KA5* | 14 | 91335086 | 91526993 | 6.66X10-05 | 7.41X10-12 | 2.74X10-14 |
| *MON1A* | 3 | 49946302 | 49967445 | 1.54X10-02 | 3.07X10-13 | 2.92X10-14 |
| *ZFPM2* | 8 | 106330917 | 106816767 | 2.09X10-03 | 1.30X10-12 | 3.04X10-14 |
| *HIVEP2* | 6 | 143072604 | 143267495 | 1.10X10-03 | 2.74X10-12 | 4.49X10-14 |
| *SLC38A3* | 3 | 50242692 | 50258406 | 2.18X10-04 | 8.20X10-12 | 6.19X10-14 |
| *TMEM161B* | 5 | 87485450 | 87564665 | 1.93X10-05 | 3.17X10-11 | 6.86X10-14 |
| *RPLP2* | 11 | 808841 | 827592 | 3.19X10-06 | 5.33X10-11 | 7.01X10-14 |
| *TCTA* | 3 | 49449639 | 49453909 | 5.22X10-04 | 9.34X10-12 | 9.80X10-14 |
| *NKAIN2* | 6 | 124124991 | 125146786 | 1.29X10-02 | 1.24X10-12 | 1.03X10-13 |
| *LONRF2* | 2 | 100889753 | 100939195 | 1.59X10-02 | 1.60X10-12 | 1.56X10-13 |
| *C14orf159* | 14 | 91526677 | 91691976 | 6.61X10-03 | 3.18X10-12 | 1.66X10-13 |
| *DUSP6* | 12 | 89741837 | 89746296 | 2.24X10-09 | 2.34X10-09 | 2.46X10-13 |
| *BACE2* | 21 | 42539728 | 42654461 | 1.26X10-03 | 1.40X10-11 | 2.63X10-13 |
| *PPP1R3A* | 7 | 113516882 | 113559082 | 1.40X10-02 | 4.29X10-12 | 3.82X10-13 |
| *APEH* | 3 | 49711427 | 49720936 | 7.59X10-03 | 6.86X10-12 | 3.92X10-13 |
| *C3orf38* | 3 | 88198875 | 88207115 | 7.37X10-03 | 7.05X10-12 | 4.02X10-13 |
| *MLIP* | 6 | 53883714 | 54131078 | 5.43X10-03 | 1.18X10-11 | 5.42X10-13 |
| *BTBD2* | 19 | 1985437 | 2015702 | 3.79X10-04 | 6.66X10-11 | 7.05X10-13 |
| *ZNF131* | 5 | 43120985 | 43176351 | 1.12X10-02 | 9.46X10-12 | 7.18X10-13 |
| *NIM1K* | 5 | 43192170 | 43280952 | 5.14X10-03 | 1.59X10-11 | 7.31X10-13 |
| *GRIA1* | 5 | 152870084 | 153193429 | 4.87X10-03 | 1.88X10-11 | 8.11X10-13 |
| *SWI5* | 9 | 131037663 | 131051268 | 1.60X10-03 | 3.82X10-11 | 8.94X10-13 |
| *DIABLO* | 12 | 122692209 | 122712081 | 1.45X10-02 | 1.01X10-11 | 9.36X10-13 |
| *PPP2R3A* | 3 | 135684515 | 135866752 | 1.04X10-02 | 1.46X10-11 | 1.07X10-12 |
| *CGGBP1* | 3 | 88101100 | 88199016 | 7.91X10-03 | 2.32X10-11 | 1.40X10-12 |
| *TEAD3* | 6 | 35441374 | 35464884 | 1.14X10-02 | 1.84X10-11 | 1.46X10-12 |
| *PPARD* | 6 | 35310335 | 35395968 | 1.38X10-02 | 1.68X10-11 | 1.52X10-12 |
| *ATP5G1* | 17 | 46970148 | 46973233 | 4.69X10-03 | 3.78X10-11 | 1.58X10-12 |
| *ARHGAP1* | 11 | 46698625 | 46722215 | 8.00X10-03 | 2.72X10-11 | 1.71X10-12 |
| *CADPS2* | 7 | 121958478 | 122526813 | 1.16X10-02 | 2.28X10-11 | 1.81X10-12 |
| *TRIM38* | 6 | 25962917 | 25987557 | 1.38X10-02 | 1.97X10-11 | 1.87X10-12 |
| *JADE2* | 5 | 133860003 | 133918918 | 9.86X10-03 | 2.71X10-11 | 1.94X10-12 |
| *LOC101929490* | 8 | 11537185 | 11555493 | 4.12X10-03 | 5.50X10-11 | 2.26X10-12 |
| *IQSEC1* | 3 | 12938542 | 13114652 | 1.12X10-03 | 1.25X10-10 | 2.47X10-12 |
| *HYAL1* | 3 | 50337320 | 50349812 | 7.66X10-05 | 4.50X10-10 | 3.27X10-12 |
| *ZNF654* | 3 | 88108394 | 88193814 | 8.17X10-03 | 5.48X10-11 | 3.46X10-12 |
| *GTF2I* | 7 | 74071991 | 74175022 | 7.30X10-03 | 6.04X10-11 | 3.59X10-12 |
| *PIDD1* | 11 | 799179 | 809872 | 5.30X10-07 | 4.17X10-09 | 4.17X10-12 |
| *ULK4* | 3 | 41288090 | 42056080 | 1.06X10-02 | 6.46X10-11 | 4.88X10-12 |
| *HIST1H2BD* | 6 | 26157419 | 26171577 | 7.23X10-03 | 9.75X10-11 | 5.98X10-12 |
| *MAP4* | 3 | 47892180 | 48130769 | 2.32X10-03 | 1.98X10-10 | 6.04X10-12 |

| | | | | | | |
|---|---|---|---|---|---|---|
| *BBX* | 3 | 107241783 | 107530176 | 2.35X10-04 | 7.90X10-10 | 7.34X10-12 |
| *BAK1* | 6 | 33540323 | 33548072 | 3.21X10-03 | 2.01X10-10 | 7.71X10-12 |
| *PNPLA2* | 11 | 818901 | 825573 | 1.18X10-04 | 1.19X10-09 | 9.37X10-12 |
| *ZFHX3* | 16 | 72816784 | 73092534 | 5.39X10-03 | 2.29X10-10 | 1.16X10-11 |
| *SLX4* | 16 | 3631182 | 3661585 | 9.87X10-03 | 1.65X10-10 | 1.22X10-11 |
| *DAG1* | 3 | 49506136 | 49573051 | 1.61X10-03 | 8.15X10-10 | 2.09X10-11 |
| *RHOA* | 3 | 49396569 | 49449526 | 5.92X10-04 | 1.80X10-09 | 2.87X10-11 |
| *PLCL1* | 2 | 198669426 | 199014608 | 3.18X10-03 | 7.85X10-10 | 2.99X10-11 |
| *TM6SF2* | 19 | 19374841 | 19384074 | 1.15X10-03 | 1.26X10-09 | 3.26X10-11 |
| *DHX30* | 3 | 47844399 | 47891686 | 5.26X10-04 | 2.22X10-09 | 3.35X10-11 |
| *SUGP1* | 19 | 19387320 | 19431321 | 3.20X10-04 | 2.53X10-09 | 3.54X10-11 |
| *ANTXR2* | 4 | 80822771 | 80994626 | 5.62X10-05 | 6.63X10-09 | 3.82X10-11 |
| *SMARCC1* | 3 | 47627378 | 47823405 | 5.56X10-04 | 2.47X10-09 | 4.11X10-11 |
| *KMT2D* | 12 | 49412758 | 49453935 | 7.34X10-04 | 2.30X10-09 | 4.33X10-11 |
| *CTNNB1* | 3 | 41236401 | 41281939 | 2.24X10-04 | 5.32X10-09 | 5.75X10-11 |
| *ICA1L* | 2 | 203637873 | 203736708 | 2.28X10-04 | 5.37X10-09 | 6.40X10-11 |
| *SCN2A* | 2 | 165986659 | 166248820 | 2.87X10-03 | 2.00X10-09 | 7.44X10-11 |
| *CSNK1G2* | 19 | 1941148 | 1981337 | 6.30X10-05 | 1.13X10-08 | 8.74X10-11 |
| *CSE1L* | 20 | 47662783 | 47713497 | 9.32X10-03 | 1.20X10-09 | 8.92X10-11 |
| *CSRNP3* | 2 | 166326157 | 166545917 | 3.81X10-03 | 2.08X10-09 | 9.06X10-11 |
| *WDPCP* | 2 | 63348518 | 63815867 | 1.28X10-03 | 4.86X10-09 | 1.21X10-10 |
| *ANKRD28* | 3 | 15708743 | 15901053 | 2.79X10-03 | 3.24X10-09 | 1.29X10-10 |
| *SLC9B2* | 4 | 103946647 | 103998480 | 1.59X10-04 | 1.29X10-08 | 1.37X10-10 |
| *SLC25A22* | 11 | 790475 | 798269 | 6.11X10-03 | 2.30X10-09 | 1.39X10-10 |
| *STK32C* | 10 | 133996038 | 134145377 | 5.33X10-03 | 2.52X10-09 | 1.42X10-10 |
| *ITPR3* | 6 | 33587951 | 33664351 | 1.81X10-04 | 1.40X10-08 | 1.45X10-10 |
| *CSPG5* | 3 | 47603728 | 47621730 | 1.55X10-03 | 5.91X10-09 | 1.68X10-10 |
| *IP6K3* | 6 | 33689415 | 33714762 | 8.75X10-04 | 9.63X10-09 | 2.05X10-10 |
| *ATP13A2* | 1 | 17312453 | 17338467 | 4.97X10-03 | 3.97X10-09 | 2.12X10-10 |
| *CDHR4* | 3 | 49828165 | 49837254 | 1.51X10-04 | 2.53X10-08 | 2.52X10-10 |
| *UQCC2* | 6 | 33664538 | 33679528 | 5.88X10-04 | 1.50X10-08 | 2.67X10-10 |
| *TOX3* | 16 | 52471682 | 52581714 | 7.43X10-03 | 4.10X10-09 | 2.73X10-10 |
| *AMBRA1* | 11 | 46417962 | 46615619 | 1.03X10-03 | 1.14X10-08 | 2.80X10-10 |
| *MANBA* | 4 | 103552643 | 103682151 | 5.99X10-08 | 4.25X10-07 | 3.70X10-10 |
| *RASGRF1* | 15 | 79252289 | 79383215 | 1.15X10-03 | 1.48X10-08 | 3.79X10-10 |
| *MFAP2* | 1 | 17300997 | 17308081 | 7.74X10-03 | 5.55X10-09 | 3.87X10-10 |
| *FOXP2* | 7 | 113726365 | 114333827 | 1.10X10-06 | 2.01X10-07 | 4.03X10-10 |
| *GLYR1* | 16 | 4853204 | 4897383 | 8.79X10-03 | 5.42X10-09 | 4.15X10-10 |
| *HIST1H4A* | 6 | 26021907 | 26022278 | 7.55X10-04 | 2.10X10-08 | 5.22X10-10 |
| *WDR12* | 2 | 203745323 | 203776949 | 2.44X10-04 | 4.18X10-08 | 5.73X10-10 |
| *BTD* | 3 | 15642864 | 15689147 | 4.72X10-03 | 1.07X10-08 | 6.11X10-10 |
| *GRIK5* | 19 | 42502468 | 42574278 | 7.58X10-03 | 9.27X10-09 | 6.48X10-10 |
| *DAGLA* | 11 | 61447905 | 61514474 | 1.99X10-05 | 1.30X10-07 | 7.09X10-10 |
| *LEMD2* | 6 | 33738990 | 33756906 | 1.33X10-03 | 2.56X10-08 | 7.21X10-10 |
| *NBEAL1* | 2 | 203879597 | 204091101 | 1.15X10-03 | 2.51X10-08 | 7.56X10-10 |
| *CHST10* | 2 | 101008322 | 101034130 | 1.36X10-02 | 8.33X10-09 | 8.25X10-10 |

| | | | | | | |
|---|---|---|---|---|---|---|
| *RMDN1* | 8 | 87479627 | 87526567 | 5.00X10-03 | 1.54X10-08 | 8.43X10-10 |
| *CALB2* | 16 | 71392616 | 71424341 | 3.91X10-03 | 1.99X10-08 | 9.69X10-10 |
| *CRIM1* | 2 | 36583370 | 36778278 | 7.34X10-03 | 1.47X10-08 | 1.01X10-9 |
| *SLC3A1* | 2 | 44502597 | 44547963 | 6.16X10-03 | 1.70X10-08 | 1.07X10-9 |
| *FOXP1* | 3 | 71003865 | 71633140 | 2.58X10-03 | 2.94X10-08 | 1.19X10-9 |
| *HMGB4* | 1 | 34326076 | 34330392 | 1.09X10-02 | 1.51X10-08 | 1.33X10-9 |
| *DGKZ* | 11 | 46354455 | 46402104 | 2.49X10-04 | 9.41X10-08 | 1.37X10-9 |
| *ATG13* | 11 | 46638826 | 46697569 | 3.07X10-03 | 3.09X10-08 | 1.39X10-9 |
| *ZKSCAN4* | 6 | 28212404 | 28227030 | 1.15X10-02 | 1.79X10-08 | 1.66X10-9 |
| *MAD1L1* | 7 | 1855428 | 2272583 | 1.58X10-03 | 5.93X10-08 | 1.93X10-9 |
| *MAU2* | 19 | 19431496 | 19469563 | 5.47X10-04 | 9.22X10-08 | 2.09X10-9 |
| *MMP24* | 20 | 33814539 | 33864804 | 1.45X10-05 | 4.13X10-07 | 2.38X10-9 |
| *PREPL* | 2 | 44544746 | 44589001 | 3.91X10-03 | 5.05X10-08 | 2.59X10-9 |
| *DDN* | 12 | 49388933 | 49393088 | 2.58X10-04 | 2.18X10-07 | 3.47X10-9 |
| *UBE2D3* | 4 | 103715540 | 103790050 | 4.22X10-05 | 4.19X10-07 | 3.52X10-9 |
| *SLC9B1* | 4 | 103806205 | 103947552 | 1.06X10-04 | 3.28X10-07 | 3.70X10-9 |
| *PDDC1* | 11 | 767222 | 777501 | 1.39X10-05 | 6.10X10-07 | 3.80X10-9 |
| *OR4C13* | 11 | 49973943 | 49974971 | 5.47X10-04 | 1.70X10-07 | 3.98X10-9 |
| *BCL2L13* | 22 | 18111621 | 18213621 | 4.67X10-03 | 7.22X10-08 | 4.14X10-9 |
| *LOC101927844* | 1 | 87678352 | 87717014 | 2.10X10-03 | 1.27X10-07 | 4.94X10-9 |
| *SIDT2* | 11 | 117049626 | 117068161 | 8.41X10-03 | 8.04X10-08 | 6.49X10-9 |
| *GATAD2A* | 19 | 19496642 | 19619741 | 4.09X10-03 | 1.07X10-07 | 6.56X10-9 |
| *RPS10* | 6 | 34385231 | 34393902 | 5.96X10-03 | 1.10X10-07 | 7.21X10-9 |
| *NICN1* | 3 | 49459766 | 49466777 | 8.41X10-05 | 6.79X10-07 | 7.21X10-9 |
| *CISD2* | 4 | 103749224 | 103813964 | 3.93X10-05 | 1.02X10-06 | 9.11X10-9 |
| *SNX14* | 6 | 86215214 | 86303850 | 8.47X10-03 | 1.11X10-07 | 9.14X10-9 |
| *PHF2* | 9 | 96338909 | 96441869 | 6.68X10-03 | 1.37X10-07 | 9.72X10-9 |
| *GRK4* | 4 | 2965232 | 3042474 | 1.41X10-03 | 3.46X10-07 | 1.19X10-8 |
| *STYX* | 14 | 53196883 | 53241707 | 4.02X10-03 | 2.18X10-07 | 1.23X10-8 |
| *RASSF1* | 3 | 50367217 | 50378367 | 1.54X10-03 | 3.94X10-07 | 1.40X10-8 |
| *GRM4* | 6 | 33989623 | 34123399 | 1.17X10-03 | 4.30X10-07 | 1.44X10-8 |
| *HYAL3* | 3 | 50330259 | 50336899 | 6.44X10-06 | 2.98X10-06 | 1.69X10-8 |
| *HPS5* | 11 | 18300217 | 18343751 | 3.27X10-03 | 3.38X10-07 | 1.74X10-8 |
| *PMFBP1* | 16 | 72152996 | 72206349 | 1.07X10-02 | 1.94X10-07 | 1.86X10-8 |
| *PKP4* | 2 | 159313476 | 159537941 | 8.49X10-04 | 6.64X10-07 | 1.95X10-8 |
| *TMEM184B* | 22 | 38612415 | 38669040 | 1.11X10-02 | 1.99X10-07 | 1.96X10-8 |
| *DIS3L* | 15 | 66585633 | 66626236 | 1.15X10-02 | 2.14X10-07 | 2.12X10-8 |
| *FAM13A* | 4 | 89647105 | 90032549 | 1.32X10-04 | 1.52X10-06 | 2.25X10-8 |
| *SYNCRIP* | 6 | 86317502 | 86353568 | 5.14X10-03 | 4.02X10-07 | 2.62X10-8 |
| *MDH1* | 2 | 63815743 | 63834331 | 2.87X10-03 | 5.83X10-07 | 2.93X10-8 |
| *BANK1* | 4 | 102711764 | 102995969 | 2.20X10-03 | 6.59X10-07 | 2.99X10-8 |
| *SDK1* | 7 | 3341080 | 4308632 | 7.22X10-03 | 4.50X10-07 | 3.60X10-8 |
| *NOP14* | 4 | 2939663 | 2965233 | 1.62X10-03 | 9.18X10-07 | 3.74X10-8 |
| *OLFM4* | 13 | 53602876 | 53626196 | 1.81X10-04 | 2.41X10-06 | 4.30X10-8 |
| *KAT2B* | 3 | 20081524 | 20195896 | 7.35X10-04 | 1.63X10-06 | 4.75X10-8 |
| *GRID2* | 4 | 93225453 | 94695707 | 2.62X10-03 | 1.04X10-06 | 5.31X10-8 |

| | | | | | | |
|---|---|---|---|---|---|---|
| *PEAK1* | 15 | 77400498 | 77712446 | 5.80X10-03 | 1.11X10-06 | 8.30X10-8 |
| *PKD1L3* | 16 | 71963441 | 72033877 | 4.85X10-03 | 1.23X10-06 | 8.97X10-8 |
| *CREB3L1* | 11 | 46299189 | 46342972 | 4.77X10-03 | 1.47X10-06 | 1.09X10-7 |
| *CREB3L3* | 19 | 4153598 | 4173051 | 8.36X10-03 | 1.22X10-06 | 1.12X10-7 |
| *MALRD1* | 10 | 19337700 | 20023407 | 2.99X10-03 | 2.18X10-06 | 1.23X10-7 |
| *THUMPD3* | 3 | 9404660 | 9428475 | 5.96X10-03 | 1.64X10-06 | 1.26X10-7 |
| *ADGRB2* | 1 | 32192718 | 32230494 | 2.96X10-03 | 2.20X10-06 | 1.27X10-7 |
| *PBXIP1* | 1 | 154916553 | 154928624 | 5.91X10-03 | 1.74X10-06 | 1.40X10-7 |
| *CD247* | 1 | 167399877 | 167487847 | 3.08X10-04 | 6.06X10-06 | 1.43X10-7 |
| *OR4C12* | 11 | 50003009 | 50004071 | 5.36X10-04 | 5.00X10-06 | 1.44X10-7 |
| *ZAK* | 2 | 173940440 | 174132737 | 9.96X10-04 | 4.09X10-06 | 1.51X10-7 |
| *PRKAG1* | 12 | 49396055 | 49413012 | 8.05X10-04 | 4.59X10-06 | 1.56X10-7 |
| *CELF4* | 18 | 34823003 | 35146000 | 1.93X10-03 | 4.00X10-06 | 1.97X10-7 |
| *TALDO1* | 11 | 747417 | 765024 | 5.64X10-05 | 1.45X10-05 | 2.07X10-7 |
| *MRPL21* | 11 | 68658744 | 68671303 | 6.73X10-03 | 3.08X10-06 | 2.73X10-7 |
| *DCC* | 18 | 49866542 | 51062273 | 5.51X10-04 | 9.61X10-06 | 3.02X10-7 |
| *ZNF521* | 18 | 22641888 | 22932214 | 1.16X10-04 | 1.91X10-05 | 3.88X10-7 |
| *CEND1* | 11 | 787110 | 790126 | 2.35X10-04 | 1.51X10-05 | 3.92X10-7 |
| *MARCH5* | 10 | 94050920 | 94113721 | 8.80X10-04 | 1.02X10-05 | 3.93X10-7 |
| *FOXO1* | 13 | 41129801 | 41240734 | 4.16X10-03 | 6.10X10-06 | 4.38X10-7 |
| *GALNT13* | 2 | 154728426 | 155310489 | 3.93X10-03 | 6.45X10-06 | 4.50X10-7 |
| *CARF* | 2 | 203776978 | 203851060 | 3.37X10-04 | 1.73X10-05 | 5.51X10-7 |
| *RNF115* | 1 | 145610990 | 145689005 | 4.22X10-04 | 2.26X10-05 | 7.08X10-7 |
| *PCSK7* | 11 | 117075788 | 117102811 | 2.23X10-03 | 1.40X10-05 | 8.45X10-7 |
| *ZNF564* | 19 | 12636184 | 12691789 | 2.08X10-04 | 3.38X10-05 | 9.83X10-7 |
| *ANO10* | 3 | 43407818 | 43663560 | 4.58X10-05 | 7.98X10-05 | 1.42X10-6 |
| *LIN28B* | 6 | 105384874 | 105531207 | 2.87X10-03 | 2.20X10-05 | 1.50X10-6 |
| *CYP20A1* | 2 | 204103164 | 204170563 | 2.23X10-03 | 2.50X10-05 | 1.72X10-6 |
| *ETF1* | 5 | 137841782 | 137878989 | 4.74X10-03 | 2.38X10-05 | 2.01X10-6 |
| *CPEB3* | 10 | 93806452 | 94050875 | 6.55X10-03 | 2.16X10-05 | 2.09X10-6 |
| *DPYSL5* | 2 | 27070969 | 27173219 | 4.56X10-03 | 2.57X10-05 | 2.16X10-6 |
| *KLHDC8B* | 3 | 49208987 | 49213919 | 2.76X10-03 | 4.02X10-05 | 2.87X10-6 |
| *UBE2J1* | 6 | 90036344 | 90062619 | 1.88X10-03 | 4.71X10-05 | 2.94X10-6 |

Gene locations are given as chromosome (CHR) and transcribed region (START and STOP sites) in the Human Genome Build 37, according to the NCBI 37.3 gene definitions.

**Supplementary Table S3.** Results of the ADHD-obesity gene-based cross-disorder meta-analysis showing the 9 genome-wide significant genes that increased significance in the meta-analysis by at least one order of magnitude compared to both individual GWASs.

| GENE | CHR | START | STOP | ADHD gene-based results | BMI gene-based results | ADHD-obesity gene-based meta-analysis |
|---|---|---|---|---|---|---|
| *BDNF* | 11 | 27676440 | 27743605 | 1.13X10-03 | 5.28X10-11 | 1.23x10-12 |
| *TFAP2B* | 6 | 50786439 | 50815326 | 1.37X10-04 | 1.20X10-08 | 1.48x10-11 |
| *DNM1* | 9 | 130965634 | 131017528 | 2.46X10-04 | 2.40X10-05 | 4.60x10-8 |
| *CACNA1D* | 3 | 53529076 | 53847179 | 3.60X10-04 | 3.64X10-05 | 9.89x10-8 |
| *BTBD2* | 19 | 1985437 | 2015702 | 3.79X10-04 | 4.47X10-05 | 1.22x10-7 |
| *FBXL17* | 5 | 107194734 | 107718080 | 4.08X10-05 | 2.61X10-04 | 1.43x10-7 |
| *SWI5* | 9 | 131037663 | 131051268 | 1.60X10-03 | 2.00X10-05 | 2.18x10-7 |
| *CSNK1G2* | 19 | 1941148 | 1981337 | 6.30X10-05 | 9.00X10-04 | 8.60x10-7 |
| *CAMKMT* | 2 | 44589043 | 44999731 | 1.63X10-04 | 6.61X10-04 | 1.17x10-6 |

Gene locations are given as chromosome (CHR) and transcribed region (START and STOP sites) in the Human Genome Build 37, according to the NCBI 37.3 gene definitions.
